# Reward gates auditory pattern encoding in the primate globus pallidus

**DOI:** 10.64898/2026.01.29.702435

**Authors:** Sujaya Neupane, Mehrdad Kashefi, Rhonda Kersten, J. Andrew Pruszynski, Jessica A. Grahn, Jonathan A. Michaels

**Affiliations:** School of Kinesiology and Health Science, York University, Toronto, Ontario, Canada; Coherent Research, coherentresearch.org, Montreal, Quebec, Canada; Western Centre for Brain and Mind, Western University, London, Ontario, Canada; Department of Physiology and Pharmacology, Western University, London, Ontario, Canada; Department of Psychology, Western University, London, Ontario, Canada; Centre for Vision Research, York University, Toronto, Canada; Centre for Integrative and Applied Neuroscience, York University, Toronto, Canada

## Abstract

The ability to anticipate rhythmic patterns is fundamental to human experience, enabling music appreciation, speech comprehension, and dancing in sync to music. How the brain learns to use acoustic information to guide motor behavior remains a key question, with its neural underpinnings and evolutionary origins debated, especially in non-human primates. Reward contingencies improve monkeys’ ability to synchronize with musical beats with near-human accuracy. However, humans and certain birds spontaneously synchronize with musical beats. Behavioural work in monkeys suggests that the brain’s reward system can encode rhythmic patterns if sufficiently reinforced, known as the *intrinsic reward hypothesis*. To investigate the neural mechanism of how the reward gates rhythmic pattern learning, we recorded neural activity from the globus pallidus interna (GPi) of a rhesus macaque during passive listening to auditory patterns. We found that GPi’s ability to encode auditory patterns in a monkey is shaped by reward association. Specifically, during passive listening (Experiment 1), with reward only at the end of each trial, GPi showed at most a weak response to the auditory tone patterns that was not evident in single neurons or population responses, ruling out spontaneous entrainment of motor activity to auditory patterns. When each tone was simultaneously accompanied by a liquid reward (Experiment 2), GPi showed clear responses to the tone patterns. To test if the tone responses persisted, the reward contingency was subsequently removed (Experiment 3). Strikingly, although the reward was delivered only at the trial’s end as in Experiment 1, GPi neurons continued to track the occurrence of the tones. Besides GPi, we also recorded single-neuron activity across the primary somatosensory (S1), primary motor (M1), dorsal premotor (PMd), supplementary motor (SMA), pre-supplementary motor (preSMA) cortices, and the medial geniculate body (MGB) of the same animal in experiments 1 and 2. We found that during passive listening (Experiment 1) with a reward only at the end of each trial, primarily the MGB, not motor areas, responded to the auditory tone patterns, ruling out spontaneous entrainment of motor activity to auditory patterns across the primate motor system. When each tone was simultaneously accompanied by a liquid reward (Experiment 2), all areas except S1 showed clear responses to the tone patterns. These findings suggest that while basic sensory processing of tone patterns occurs in the MGB, the GPi exhibits reward-driven encoding of auditory patterns that persists after reward withdrawal. This reorganization of neural responsiveness shows that the capacity for auditory association can be elicited in the motor system through appropriate motivational contexts.

## Introduction

Humans have a remarkable ability to extract rhythmic structure from sound and to align perceptual expectations and movement around a steady beat. From spontaneous dancing to ensemble performance, people synchronize actions with temporal precision across a broad range of tempi and contexts^1,2^. This capacity for beat perception and synchronization relies on tight coupling between auditory and motor systems and recruits a distributed network spanning auditory cortex, supplementary and premotor areas, basal ganglia, and cerebellum^3–7^. Reward and affective circuits are integral to this system: activity in the caudate and ventral striatum tracks both musical pleasure and rhythmic engagement, suggesting that timing, prediction, and motivation are deeply intertwined^4^.

Whether such rhythmic capacities are uniquely human remains debated. The *vocal learning hypothesis* proposes that only species capable of complex vocal mimicry evolved the auditory-motor coupling necessary for beat perception and synchronization^8,9^. The *gradual audiomotor evolution hypothesis* instead argues for continuity: interval-based timing is broadly conserved across primates^10–13^ and rodents^14–16^, whereas hierarchical beat perception represents an extension within humans^17^. Rhesus macaques exemplify this continuum. They produce precise temporal intervals^12,18^, detect rhythmic groupings without perceiving a beat^19^, and can learn to synchronize taps to metronomes or adapt to changing tempi, though their movements often lag the beat rather than anticipating it^20,21^. These findings imply that primates possess the requisite neural machinery for rhythmic timing but that its expression may depend on sensory modality and motivational state.

Neurophysiological evidence situates rhythm within distributed cortico-basal ganglia-thalamic loops. Medial premotor neurons are tuned to interval duration during isochronous tapping^5^, parietal populations ramp toward internally generated movement onsets^12^, and predictive dynamics emerge in motor thalamus and cerebellum^22–24^. In humans, these same circuits overlap with networks mediating attention, arousal, and reward^4,25^, consistent with the idea that motivational engagement modulates access to timing mechanisms.

Recent work reinforces this link between motivation and rhythmic behavior. Reward contingencies improve monkeys’ synchronization precision^20,21^, and under rich reinforcement schedules, they can even align to musical beats with near-human accuracy^26^. These results suggest that the expression of rhythmic ability depends less on sensorimotor limitations than on whether rhythmic sounds acquire motivational value, leading to the proposal of the *intrinsic reward hypothesis*^*20*^. Under this hypothesis, learning to perform synchronized movements to auditory rhythm should first engage the brain’s reward circuitry to associate patterned sound with its intrinsic value, even in the absence of movement. Yet, nearly all existing studies in non-human primates require overt movement or explicit prediction, leaving unresolved whether auditory patterns alone, in the absence of any task demand, can recruit reward and sensorimotor circuits for encoding auditory temporal structure^27^. In humans, even passive listening to rhythmic stimuli activates motor areas^28^, whereas this question has not been investigated in other primates.

To test this, we recorded neural activity from the output node of the basal ganglia, the globus pallidus interna (GPi), in a macaque that had never been trained to use auditory stimuli for behavior, while manipulating reward context but requiring no movement or prediction. In Experiment 1, where tone sequences were passively presented while reward was delivered only at trial completion, GPi along with motor areas generally did not respond to the presence of tones. In Experiment 2, each tone was paired with liquid reward, coupling auditory events to primary reinforcement. In Experiment 3, rewards were again delivered only at the end of the trial to probe the persistence of any learned representations. Although Experiments 1 and 3 were identical in terms of sensory input and motor output, the interposed Experiment 2 allowed us to test if reward drives the emergence and maintenance of auditory pattern representations in GPi. We predicted that basal ganglia areas, based on human neuroimaging evidence of modulation by pleasurable rhythm^4^ – would acquire and retain auditory responsiveness through coupling with reward. The results confirmed this prediction, supporting the *intrinsic reward hypothesis* of auditory tone perception: motivational state alone can gate the presence of auditory pattern representations in the GPi. Rather than viewing auditory pattern sensitivity as a uniquely human adaptation tied to vocal learning, our results suggest that auditory pattern responsiveness in neural circuits can arise from mechanisms that couple auditory experience to motivational value.

## Results

We used high-density Neuropixels probes to record neural activity in thalamic, basal ganglia, and cortical areas of the macaque brain during passive listening to patterned auditory stimuli. These auditory patterns, each lasting 7.3 seconds, were interleaved randomly and were either regular, irregular, or regular with every fourth tone omitted and spanned a range of tempi, allowing us to dissociate neural responses to auditory events from elapsed-time and reward-related dynamics (Fig. 1, Methods). Each tone was a high-frequency pure tone (1000 Hz) and lasted 50ms. The animal was required to maintain hand fixation and in Experiment 1 a liquid reward was given at the end of each pattern, marking the end of a trial. This reward scheme was altered in Experiment 2 and restored in Experiment 3. The animal had never been trained to perform tasks based on auditory cues, and was exposed to this task only for a few experimental sessions before recordings took place (see Supplementary Table for a summary of training and recording sessions).

**Figure 1.**
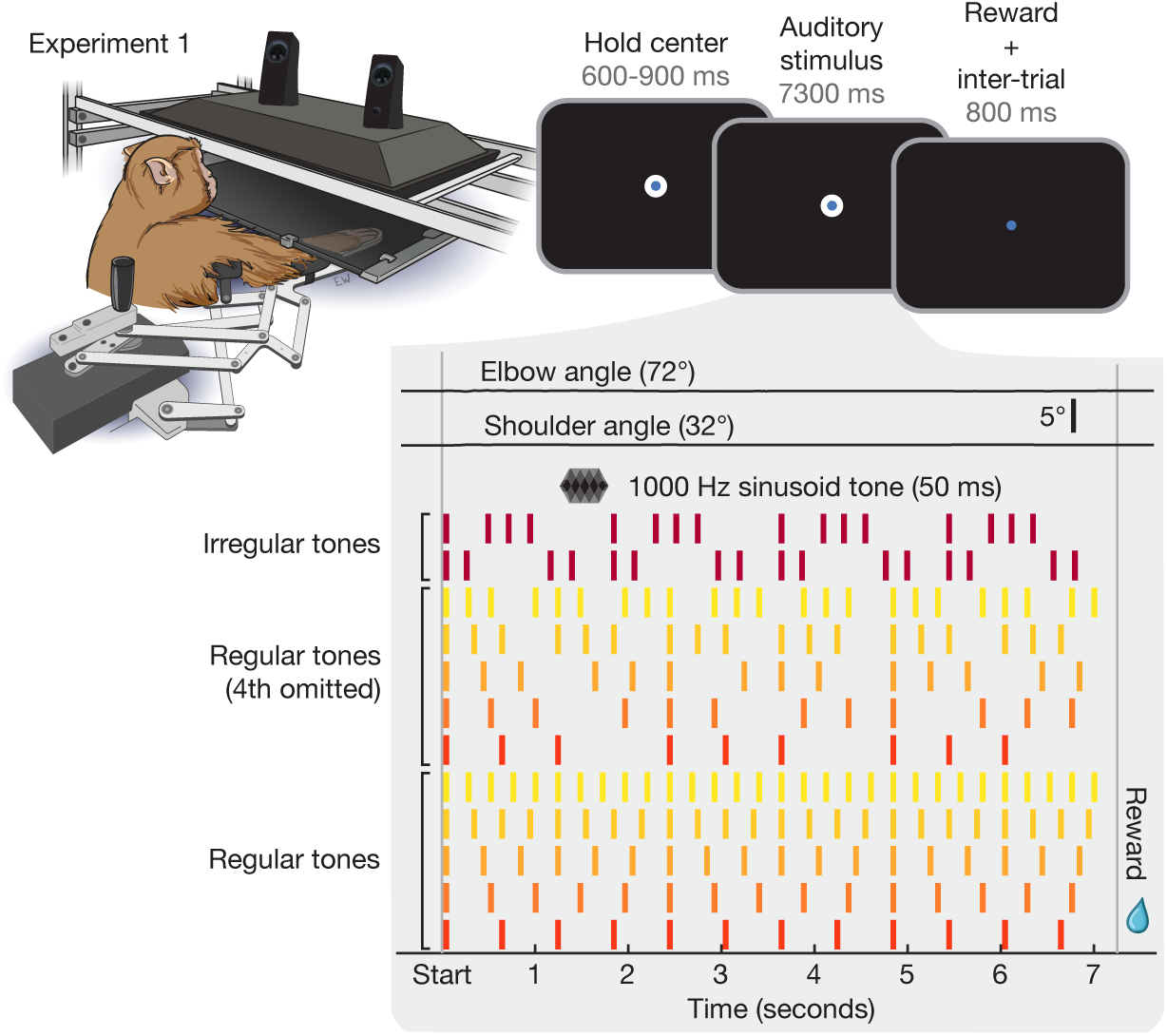
Experimental setup and paradigm. **Left**, Monkey sat in a KINARM NHP robotic exoskeleton (an arm-control setup that allows experimental manipulation of hand movement) and held the hand in place at a central location on the screen throughout the trial. **Right, top**, The experimental trials consisted of hand fixation followed by an auditory pattern stimulus and a liquid reward. **Bottom**, The animal kept the arm still throughout the time when auditory patterns were playing (example kinematics from experimental trial shown at top). Below are all auditory patterns played, from which a random one of these patterns was selected on each trial. Each tone consisted of a 50 ms pulse of a 1000 Hz sinusoid.

### Absence of entrainment to auditory patterns in the primate motor areas

In Experiment 1, we recorded activity from primary somatosensory (S1), primary motor (M1), dorsal premotor (PMd), supplementary motor (SMA), pre-supplementary motor (preSMA) cortices, globus pallidus interna (GPi), and medial geniculate body (MGB) using Neuropixels probes (Methods) while the animal passively listened to tones of different frequencies and regularities (Fig. 2a). We first examined whether neural activity across recorded areas reflected the temporal structure of the auditory sequences at the level of single neurons and population responses. We found single neurons in area MGB (but not in any other area) that showed reliable tone-locked responses (Fig. 2b right panel). However, we observed that neurons in all areas exhibited ramping activity that appeared to track the time to reward delivery at trial end (Fig. 2b left panel and Fig. S1).

**Figure 2.**
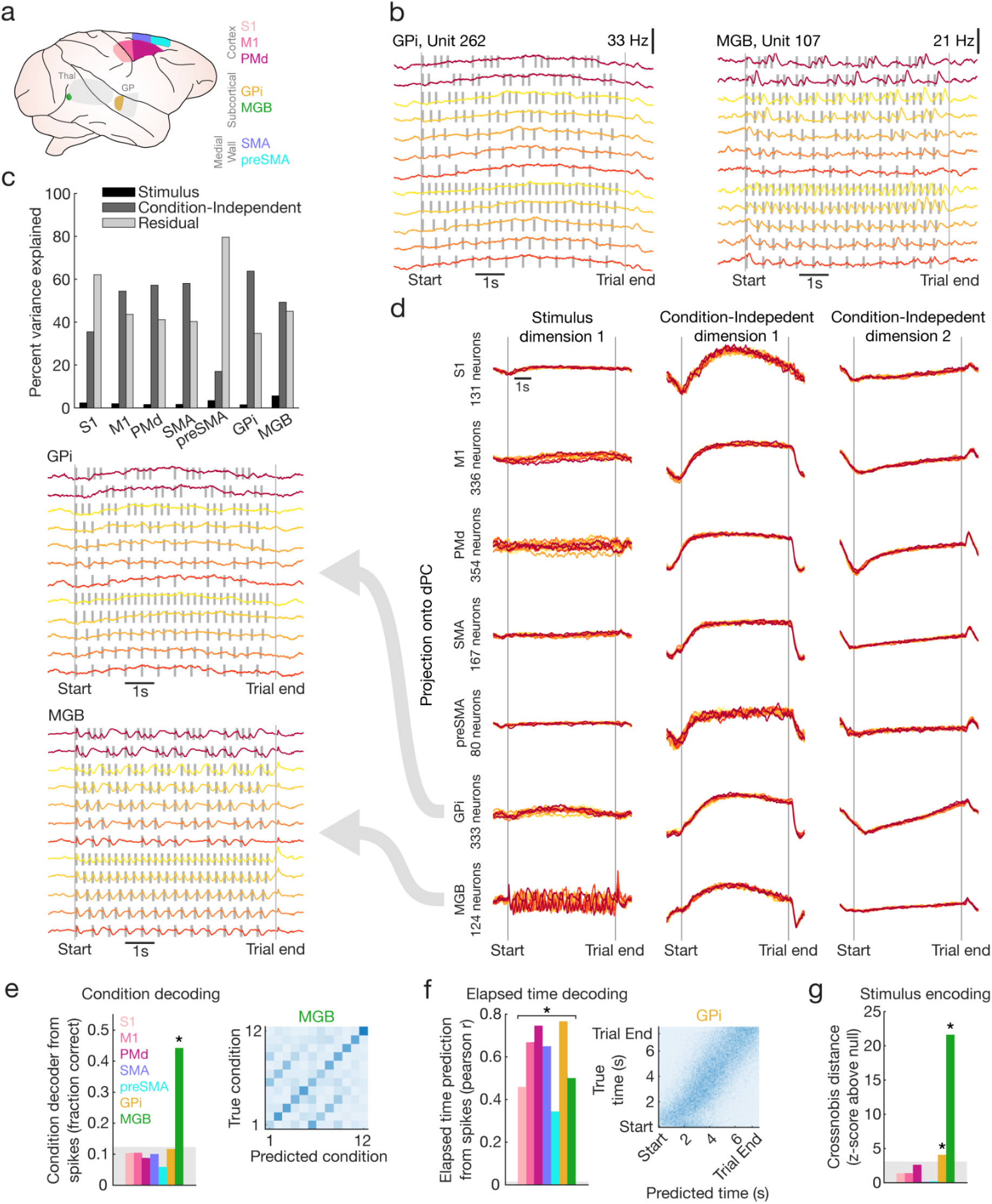
Absence of entrainment to auditory patterns in the primate motor areas. **a**, The brain areas where we recorded neural activity are color-labeled. **b**, Example single neurons from GPi (left) and MGB (right) showing modulation to timing and auditory tones, respectively. **c, Top**, Variance explained by demixed principal component (dPC) factors that represent the auditory pattern (black), condition-independent task modulation (dark grey) and residuals (light grey) across multiple brain areas. **d**, Time course of first condition-dependent and -independent dPCs across the same brain areas. **Left**, Condition-separated time course of stimulus-encoding dPC estimated from neural population activity in GPi and MGB. **e, Left**, Average stimulus-condition decoding accuracy across different brain areas. Star indicates p < 0.001 relative to chance level. **Right**, Accuracy of the condition decoder across all conditions in MGB. **f, Left**, Average elapsed time decoding accuracy across different brain areas. Star indicates p < 0.001 relative to chance level. **Right**, Accuracy of elapsed time decoder across all timepoints in GPi. **g**, Auditory pattern encoding quantified by crossnobis distance between odd-even splits of neural population activity weighted by normalized stimulus pattern. Stars indicate p < 0.001 relative to chance level.

To characterize stimulus encoding and timing signals, we employed complementary population analysis methods. We used demixed principal components analysis (dPCA, Methods) to separate condition-dependent and condition-independent components in the neural population. We reasoned that the reward tracking signal should be recovered by the condition-independent component. In general, we found that the condition-independent component explained over 50% of the variance of neural activity across all brain regions under study (Fig. 2c), except pre-SMA and to some extent S1. The main condition-independent signals visually appeared as various kinds of ramps (Fig. 2d), consistent with an event-related timer. A relatively small portion of the variance (<10%) was explained by the stimulus condition across brain regions. However, as expected from single-neuron activity, the condition-dependent component of the MGB exhibited periodic patterns, indicating stimulus encoding.

Another way to quantify how reliably neural populations tracked different auditory patterns is to test if we can decode what pattern was presented on each trial. Using a cross-validated linear decoder (Methods), we found that the decoded stimulus condition was not above chance in all brain regions except MGB (Fig. 2e), and that most of the misclassifications occurred between trials of the same frequency where the 4th tone was omitted (Fig. 2e, off-diagonal stripes in the right panel). This confusion is consistent with tone responses being purely reactive to tone onsets: even in MGB we observed no response time-locked to the omitted fourth tone, indicating the absence of any predictive or omission-related signal during passive listening. In contrast, all areas showed significant prediction of elapsed time in each trial, especially PMd and GPi (Fig. 2f, Methods).

To quantify the strength of stimulus encoding, we used crossnobis distance to evaluate how strongly auditory stimulus timing is represented relative to chance (Fig. 2g, Methods), showing that by far the strongest representation was in MGB (>20x standard deviations above null). Interestingly, there was also a significant representation in GPi (∼4 standard deviations above null), suggesting a weak representation in GPi that was not visually detectable in the population response using dPCA. No areas other than MGB and GPi showed significant stimulus encoding. These analyses (Fig. 2e-g) were only possible because of the large populations of neurons recorded simultaneously on single trials, an advantage of our high-density Neuropixels recordings.

Finally, as a complementary analysis to spiking signals, we repeated our population measures on local field potential (LFP) signals. We found that the LFP analysis generally recapitulated what we observed in spiking activity (Fig. S2), although weaker overall especially for condition decoding, indicating that the network-level representations of LFPs parallel those of the spiking representations. One notable exception was the absence of significant stimulus encoding in the LFP of GPi.

Taken together, the activity pattern across brain regions in Experiment 1 suggests that the animal was mainly tracking the underlying reward-timing signal evolving over several seconds, whereas the neural representation of the auditory pattern was limited to the activity of MGB, to a small extent GPi, and was absent in the canonical sensorimotor circuits during passive listening.

### Reward drives the emergence of auditory pattern encoding in motor circuits

The absence of robust auditory pattern encoding in motor circuits suggests that auditory pattern information is not generally available to the motor system of macaques when they are not trained to perform a task leveraging this information. This is in contrast to findings in the human brain, where motor areas are activated by passive listening to musical beats^28^. Under the *intrinsic reward hypothesis*^*20*^, synchronization to an auditory pattern occurs only if the tones are intrinsically rewarding, as is the case for humans and songbirds, whose vocal learning systems confer intrinsic value to structured auditory patterns. A key difference between humans and macaques may be the intrinsically rewarding potential of a musical piece. We therefore reasoned that pairing each tone with an explicit reward could render the auditory pattern intrinsically rewarding.

To test this idea, we conducted a second experiment in which a drop of juice reward was delivered simultaneously with each tone of the auditory patterns (Fig. 3a). We used a subset of the stimuli from Experiment 1 to maximize the number of trials per condition. We again recorded neural activity from multiple brain areas while the monkey passively listened to the auditory patterns and maintained hand fixation. Strikingly, we now observed that not only MGB, but also other brain areas – especially GPi and PMd – showed strong responses to the pattern. Single neurons in GPi and PMd were strongly modulated (Fig. 3b and Fig. S3) and showed a clear departure from the stimulus-insensitive activity observed in the first experiment when rewards were delivered at the end of a trial. Next, we applied our population analyses and found that MGB, GPi, PMd, and to a lesser extent, SMA showed robust modulation in the dPC dimension for stimulus, whereas the ramp-like modulation of the condition-independent signal was almost completely abolished across areas, with the exception of GPi (Fig. 3c,d). Decoding of the stimulus condition was above chance in all areas except S1 and pre-SMA (Fig. 3e), with the strongest decoding in PMd, GPi, and MGB. Consistent with the reduction in condition-independent signals, decoding of elapsed time was reduced across all areas except GPi (Fig. 3f). Finally, stimulus encoding was strong in many areas, except S1 where it did not reach statistical significance, and in SMA and pre-SMA where it remained at a low, but significant level (Fig. 3g).

**Figure 3.**
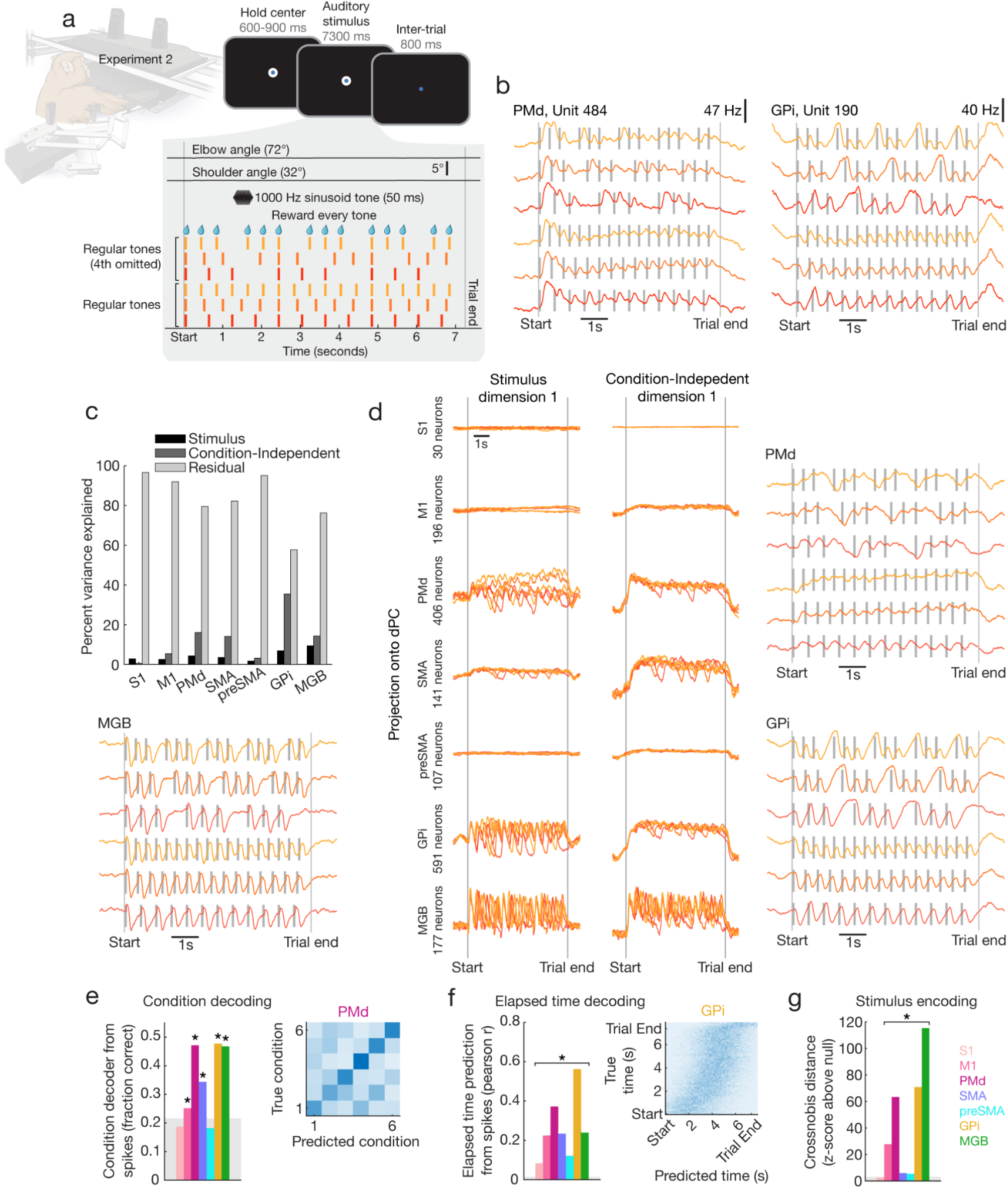
Reward drives the emergence of auditory pattern encoding in motor circuits. **a**, Experiment 2 in which each beat of the auditory pattern was paired with a liquid reward. **b**, Example single neurons from PMd (left) and GPi (right) showing modulation to auditory tones, unlike in Experiment 1. **c**, Variance explained by demixed principal component (dPC) factors that represent the auditory pattern (black), condition-independent task modulation (dark grey) and residuals (light grey) across multiple brain areas. **d**, Time course of first condition-dependent and -independent dPCs across the same brain areas. **Sides**, Condition-separated time course of stimulus-encoding dPC estimated from neural population activity in MGB, PMd, and GPi. **e, Left**, Average stimulus-condition decoding accuracy across different brain areas. Stars indicate p < 0.001 relative to chance level. **Right**, Accuracy of condition decoder across all conditions in PMd. **f, Left**, Average elapsed time decoding accuracy across different brain areas. Star indicates p < 0.001 relative to chance level. **Right**, Accuracy of elapsed time decoder across all timepoints in GPi. **g**, Auditory pattern encoding quantified by crossnobis distance between odd-even splits of neural population activity weighted by normalized stimulus pattern. Stars indicate p < 0.001 relative to chance level.

Similar to Experiment 1, LFP analysis recapitulated what we observed in spiking activity (Fig. S4), except for weaker overall decoding of stimulus patterns and no significant stimulus encoding in the LFP of GPi.

Consistent with the *intrinsic reward hypothesis*, we observed that when an auditory tone was associated with a reward, neural circuits in the macaque brain robustly represented the timing of these events. As in Experiment 1, however, these responses remained reactive rather than predictive: no area showed a response time-locked to the omitted fourth tone, suggesting that predictive, omission-sensitive responses would likely require explicit behavioural training. Is it possible that these signatures were purely driven by the motor intention to lick every time a reward was presented? To test this possibility, we used an infrared camera to record facial movements near the reward spout (Fig. S5a). We tested whether or not the stimulus condition could be decoded from face movement energy alone. For all Experiment 2 sessions, the stimulus condition was decodable from face motion energy (Fig. S5b, p < 0.001). Because each tone was paired with a reward, these orofacial movements are an expected consequence of reward delivery rather than a confound to be explained away, and they may well contribute to the responses observed in Experiment 2. Critically, this is precisely the ambiguity that Experiment 3 was designed to resolve, by testing whether tone encoding persists once the per-tone reward, and the movements that accompany it, are removed. A key question was – now that the extrinsic rewards have been paired with auditory tones and that association presumably encoded by the reward circuitries, will auditory tones be intrinsically rewarding without a paired juice reward?

### Reward-driven auditory pattern encoding in the basal ganglia persists after reward withdrawal

To test if auditory tones would now elicit responses in the basal ganglia even without extrinsic rewards, we recorded neural activity in GPi while repeating the design of Experiment 1 with the same subset of conditions from Experiment 2, once again delivering a reward only at the end of each trial (Experiment 3, Fig. 4a). The *intrinsic reward hypothesis* predicts that, after establishing an association between auditory tones and reward, the brain should maintain this association, rendering the auditory stimulus ‘rewarding’^20^. Consistent with our prediction, we saw clear representations of auditory timing in GPi in single neurons (Fig. 4b) as well as the population (Fig. 4c), in contrast to Experiment 1 – despite being identical in terms of input and task structure. Condition-independent signals and residuals were once again a dominant factor (variance explained for stimulus dimensions: 1.4%, condition-independent: 30.1%, residual: 68.5%). Consistent with these observations, the stimulus condition was now decodable from GPi to a limited extent (Fig. 4d), and crucially the stimulus encoding as measured by crossnobis distance was significant and ∼4x higher than in Experiment 1 (Fig. 4e), although notably much lower than Experiment 2. Interestingly, and similar to Experiment 1 and 2, we did not observe any significant stimulus encoding in the LFP of GPi (Fig. S6).

**Figure 4.**
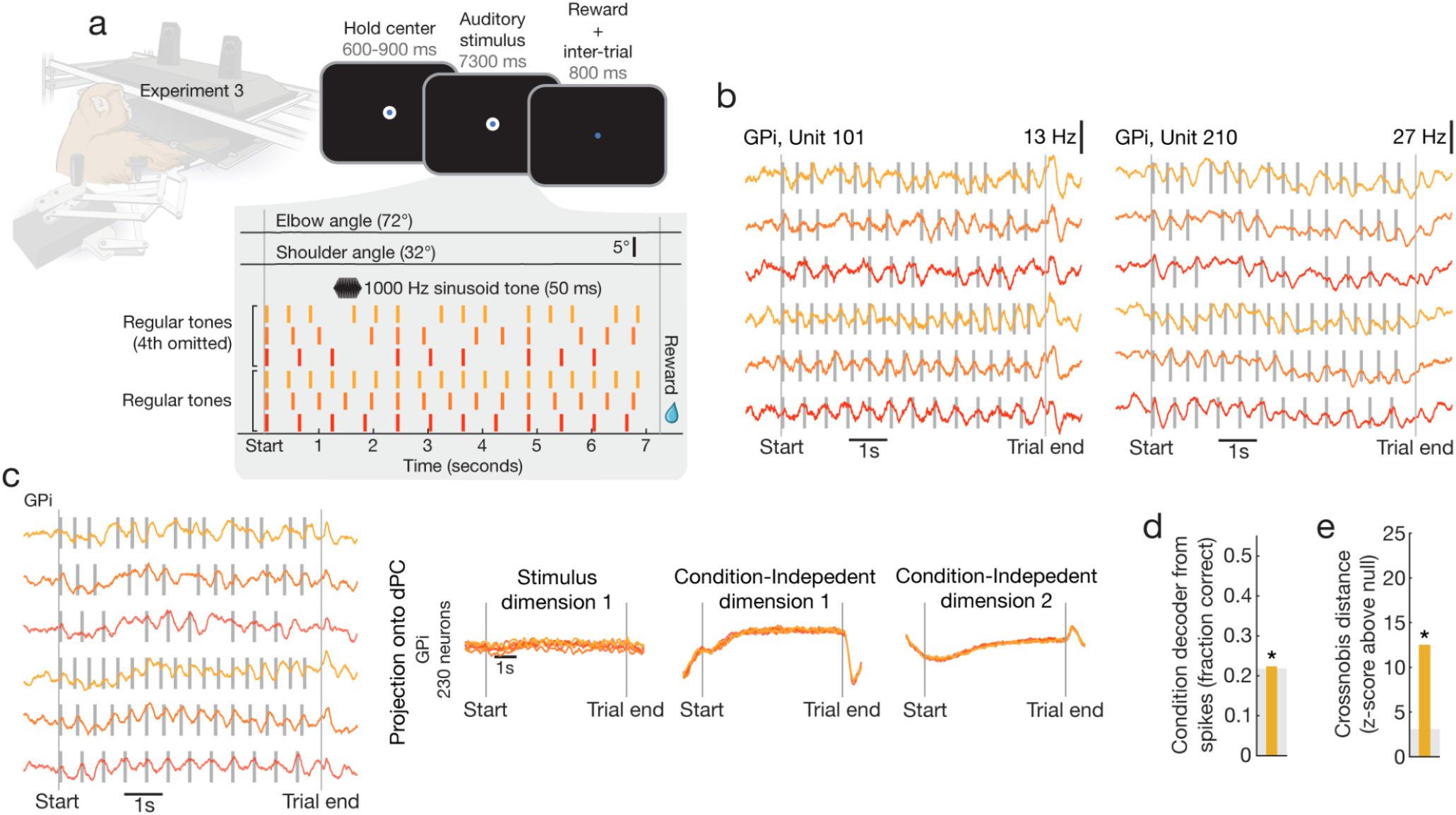
Reward-driven auditory pattern encoding in the basal ganglia persists after reward withdrawal. **a**, Experiment 3 in which reward was only delivered at the end of each trial and each tone of the auditory patterns was not paired with any reward (identical to Experiment 1). **b**, Two example single neurons from GPi showing modulation to auditory tones, similar to Experiment 2, but unlike Experiment 1. **c, Right**, Time course of condition-dependent and -independent dPCs in GPi. **Left**, Condition separated time course of stimulus dPC in GPi. **d**, Average stimulus condition decoding accuracy in GPi. Star indicates p < 0.001 relative to chance level. **e**. Auditory pattern encoding quantified by crossnobis distance in GPi. Star indicates p < 0.001 relative to chance level.

One question is how long auditory tones remain intrinsically rewarding when not coupled to behavior. This coupling may degrade rapidly when tones are no longer paired with an external reward. To test this, we compared the stimulus-encoding strength between early and late parts of Experiment 3 and found no reduction in stimulus-encoding strength, with a crossnobis distance of 5.6 in the first 250 trials and 6.6 in the last 250 trials (both lower than the full-session estimate in Fig. 4e, as crossnobis distance grows with the number of trials per split and so should not be compared directly against the full-session values reported for the other experiments). As a final control, an analysis of infrared video during Experiment 3 showed that, in contrast to Experiment 2, the stimulus condition could not be decoded from face motion energy (Fig. S5b, p = 0.76), confirming that the results of Experiment 3 cannot be explained by movement.

Together, our three experimental manipulations provide compelling empirical support for the *intrinsic reward hypothesis*. More broadly, our findings suggest that associating salient rewards with sensory inputs can unmask the basal ganglia circuits’ latent capacity to encode temporal patterns in a non-vocal-learning animal, such as a non-human primate.

## Discussion

Our results demonstrate that while the macaque motor system does not spontaneously encode auditory patterns during passive listening, the association of sound with reward can rapidly recruit one node of these circuits, the globus pallidus interna (GPi), to track rhythmic structure. In Experiment 1, we observed that neural populations across the motor system and the GPi primarily encoded elapsed time within a trial rather than auditory patterns. However, once auditory tones were paired with liquid reward in Experiment 2, a profound reorganization occurred – ramping activity diminished, and robust encoding of the auditory pattern emerged across most areas. Most strikingly, in Experiment 3, the GPi maintained this pattern encoding even after the immediate reward was withdrawn. These findings provide neurophysiological evidence for the *intrinsic reward hypothesis*^20,29^, suggesting that the neural machinery for rhythmic processing is present in non-human primates but remains latent until unmasked by motivational significance.

A central debate in the evolution of music is why humans spontaneously entrain to auditory tones. In humans, passive listening to rhythms automatically recruits motor regions^3,28^, including the premotor cortex and basal ganglia. In contrast, our results in Experiment 1 confirm that macaque motor circuits do not automatically track auditory patterns. This is consistent with behavioral studies^18^, which indicated that while macaques can quantify single intervals similarly to humans, they lack the spontaneous negative mean asynchrony (predictive timing) characteristic of human entrainment. However, the absence of spontaneous entrainment does not imply an absence of capacity. Takeya et al.^20^ demonstrated that when every predictive saccade was reinforced with immediate reward, monkeys could synchronize to a visual metronome and generalize to novel speeds. Similarly, Rajendran et al.^26^ recently demonstrated that macaques can indeed perceive and tap to a subjective beat in real music when the behavior is reinforced. Our physiological data provide a neural mechanism for these behavioral observations: the latent capacity of the basal ganglia and premotor cortex to encode auditory patterns is gated by reward.

The emergence of pattern encoding specifically in the GPi during and after reward association highlights the basal ganglia’s pivotal role in integrating timing and motivation. The putamen is critical for encoding the serial order and duration of intervals during rhythmic tapping^30^, and the striatum is a key substrate for beat processing^31^. Our finding that the GPi retains the rhythmic pattern code in Experiment 3 suggests that reward establishes a persistent ‘valuation’ of the auditory temporal structure. We propose that the GPi acts as an interface where the ‘extrinsic’ reward of the juice becomes an ‘intrinsic’ value assigned to the auditory pattern, effectively simulating the neural state of a vocal learner who finds the rhythm itself rewarding^32^.

The shift from ramping activity (time-to-reward) to tone-aligned activity (tracking the pattern) in our data reflects the flexibility of the primate motor system. Medial premotor cortex populations can represent time through circular neural trajectories that encode tempo^7,33^. Our results suggest that this sophisticated machinery, often studied in the context of motor production, can be co-opted for sensory representation when motivationally salient. In Experiment 1, the ramping activity acted as a ‘timer’ that tracked the trial duration (likely for the final reward), consistent with the evidence that the cerebellum and basal ganglia encode expectations of sensory events^22,34^. In Experiments 2 and 3, this machinery was repurposed to track the fine structure of the auditory pattern.

One interesting caveat to our findings is that we found no evidence of predictive synchronization to individual tones, even during tone omissions. Across all experiments tone responses were reactive to the presentation of the stimulus. Our observation of the emergence of stimulus encoding after reward association offers a more complex picture of how non-vocal-learning animals might learn to synchronize to rhythms. Since the underlying neural circuit for rhythmic synchronization appears to be latent in non-vocal-learning animals like macaques, learning rhythmic synchronization may not be spontaneous as in humans and parrots, but rather more gradual^20,26,35^. Although we didn’t test the behavior of the animal here, our work provides a compelling initial step for future work to test how auditory pattern responses transition from reactive to predictive.

Another caveat to our findings is that all results come from a single animal. However, by recording across three reward contingencies in the same animal, we are able to make stronger within-subject claims than between-animal designs typically afford, at the cost of generalization across individuals. Future work will test how well our within-subject claims can be replicated and extended.

While extensive, the brain regions we recorded from do not cover all regions that may be involved in learning auditory patterns. For example, we did not record from the auditory cortex and therefore we do not know if it simply follows its thalamic input (MGB), or interacts with the striatal and premotor circuits to learn auditory patterns. We also don’t know if the intrinsic reward coupling observed here is a general mechanism independent of modality or specific to the auditory and sensorimotor system, nor how quickly these associations are made and decay. Future studies combining behavioural observation and multi-region neural recordings will help answer these questions and shed light on how reward circuits interact with sensorimotor circuits to enable rhythmic entrainment.

Another question is how long the unreinforced pattern encoding in GPi persists after withdrawal of the external reward in our tone-synchronized reward scheme. We know that the pattern encoding in Experiment 3 was just as strong in the second half of the recording session as in the first half, all of which took place three days after Experiment 2 when tones were synchronized with rewards. One possibility is that once activated, the tone-reward association circuit is formed and lasts throughout the lifetime. The theory of neural recycling would suggest such a scenario^36^. According to this theory, a neural precursor evolved for one function is repurposed for new, culturally relevant functions contingent on context, such as the emergence of brain areas dedicated to recognizing letters^37,38^, performing arithmetic^39,40^, and adopting tool use^41^. Although it is challenging to test this idea in neural activity, monkeys in captivity can be behaviorally tested for this theory through a longitudinal study. Alternatively, the pattern encoding may gradually decay over time, suggesting that it is a form of memory formation and decay rather than the activation of a neural computation through recycling.

In conclusion, we show that parts of the macaque motor system, namely the GPi, are not deaf to auditory rhythm, but rather indifferent to it until recruited by reward. This reward-driven emergence of auditory pattern encoding in the motor system bridges the gap between the rigid vocal learning distinction and the behavioral flexibility observed in recent animal studies^20,26^.

## Methods

### Subject

One male rhesus macaque (Monkey P, Macaca mulatta, 16 kg, 12 years old) participated in the study. All procedures described below were approved by the Institutional Animal Care and Use Committee at Western University (Protocol #2022-0208).

### Apparatus

The subject was seated with its right arm in a KINARM NHP robot exoskeleton (Fig. 1a, BKIN Technologies^42^), allowing flexion and extension movement of the shoulder and elbow joints in the horizontal plane. The two segments of the exoskeleton, consisting of the upper arm and forearm, have adjustable cuff sizes to match the dimensions of the subject’s arm. The robot was calibrated to align a real-time cursor on the right index fingertip. The hand-position feedback and visual targets of the experiment were displayed in the same horizontal plane as the arm movement. These virtual-reality images were projected at eye-level via an LCD monitor onto a semi-silvered mirror. Before initiating the experiment, an opaque blinder was installed beneath the mirror to occlude direct vision of the physical arm during all trials. Kinematic data were sampled at 4000 Hz. For delivery of auditory stimuli, two standard computer speakers were positioned above the workspace equidistant from the subject (Fig. 1a), emitting tones at a standard volume (∼70 dB).

During Experiments 2 and 3, infrared video was recorded of the face near the reward spout using a Teledyne Genie Nano near-infrared camera (TEL-G3-GM12-M0640, recorded with a resolution of 640 × 480 pixels at approximately 60 Hz, Fig. S5).

### Experimental procedure

Our experiments involved a non-human primate passively listening to patterned auditory tones with no task-relevant behavioral demand except hand fixation. On each trial, the monkey waited with its fingertip in a central target (located under the fingertip when the shoulder and elbow angles were 32° and 72°, respectively; target size: 1.5 cm diameter). After a variable delay (600-900 ms), a 7.3 seconds-long auditory stimulus was played on two loudspeakers. Specifically, the time intervals between pulses were 0.6, 0.480, 0.4, 0.3, and 0.240 seconds for the regular tones, the same again for the regular tones but with every fourth tone omitted, and 0.225 seconds for the irregular tones, where tones were played either on the 1st, 2nd, 6th, and 7th beats of an 8 beat phrase or on the 1st, 3rd, 4th, and 5th beats of an 8 beat phrase. All tones were 1000-Hz pure tones, 50 ms in duration, with 10 ms linear rise and fall ramps. If at any point during the trial (∼8 seconds) the hand went outside the target or hand velocity within the target exceeded 0.5cm/s, the trial was aborted and rendered unsuccessful. Otherwise, the animal was rewarded with a juice reward at the end of the trial. The time between each trial (beginning of trial-end reward and the next trial start) was 800ms. The different patterns of tones are depicted in Figures 1, 3 and 4. We ran a total of 10 experimental sessions whose timeline and experimental details are shown in Table S1.

The three kinds of experimental sessions differed on how the reward was delivered with respect to the auditory stimulus patterns. In Experiments 1 and 3, the reward was only delivered at the end of the trial if the animal maintained its hand fixation. In Experiment 2, each tone of the auditory pattern in a given trial was paired simultaneously with a smaller liquid reward and no reward at the end of the trial.

### Electrophysiological recording

We performed high-density Neuropixels probe recordings (1.0 NHP - 4.5 cm). After training on basic motor tasks unrelated to this experiment, the monkey was implanted with a custom 3D-printed titanium implant (accurate to 0.2 mm) that was designed to precisely conform to its individual skull as determined by a model obtained using micro-CT. Titanium implants were fixed in the skull using a variable number of titanium screws and included a built-in recording chamber and head post. Neural recording targets were identified by registering the CT to a pre-surgery MRI, and identifying the 3D location of each brain area by warping segmentations from a composite macaque atlas to the individual MRI (NMT v2^43,44^, CHARM^43,45^ and SARM^46^ atlases, see Hirai & Jones^47^ for additional thalamic parcellations). The use of a skull conforming titanium implant allowed us to precisely plan recording trajectories to target desired structures. The precision of our implantation technique has been confirmed post-mortem in another animal to be accurate to within <0.5 mm on the cortical surface. Our method allowed insertion through the dura using 9 mm retractable guide tubes and actuated using a manual microdrive to record through small 2.7 mm craniotomies. For each recording configuration, we 3D printed a custom holder (Formlabs 3B+, Grey resin V4) that aligned the Neuropixels along a specific, pre-defined trajectory targeting the areas of interest. Recordings in the primary somatosensory cortex (S1) primarily targeted Brodmann area 3b. Recordings in M1 targeted gyral M1.

### Neural data processing

Neural data were recorded from Neuropixels probes using SpikeGLX. Drift was minimal due to small craniotomies (drift 0-15 um), so we immediately processed the data using Kilosort 4.0^48^ including built-in drift correction. Single neurons were considered successfully recorded if they were flagged by Kilosort as single neurons using default parameters and if they were stably recorded. To determine whether neurons were properly isolated over the course of the recording, we counted the number of spikes for each neuron in 30 second chunks spanning each entire recording and calculated the index of dispersion for each neuron (variance over time block divided by mean over time block). It’s important to note that this metric does not test neurons for tuning to the task, only for reliable responses over time. Neurons with an index of dispersion below 2 were included in further analysis. The majority of neurons had an index of dispersion <1, and shifting this threshold ±1 did not affect results. For the vast majority of analyses, single-unit spike trains were converted to smoothed firing-rate estimates by convolution with a causal Gaussian kernel (σ = 100 ms) prior to analysis.

Local Field Potentials (LFPs) were read from the Neuropixels LF stream, which was recorded at 2,500 Hz. All LFP processing was performed using the FieldTrip toolbox^49^. During processing the 384 channels were spatially downsampled by sampling every 12th channel, leading to a total of 32 channels. LFP data were demeaned, and a bandpass filter was applied (1-400 Hz, 3rd order). Time-frequency analysis was performed using the multi-taper convolution method with a Hanning taper and subsequently output as a power spectrum. Fifty frequencies of interest were logarithmically (log10) spaced from 1 Hz to 400 Hz. The time windows for the convolution were dynamically adjusted relative to the frequencies of interest to cover 5 cycles at each frequency. The time bins of interest were sampled at a resolution of 0.01 seconds.

### Demixed principal components analysis

Principal component analysis (PCA) is commonly employed to reduce the dimensionality of high dimensional datasets by finding a low dimensional representation that captures large amounts of variance using independent linear combinations of neurons. For PCA, given a matrix of data *X*, where each row contains the average firing rates of one neuron for all times and task conditions, PCA finds an encoder, *F*, and an equivalent decoder, *D*, which minimizes the loss function ℒ = ∥*X* − *FDX*∥ ^2^ under the constraint that the principal axes are normalized and orthogonal, and therefore *D* = *F*^*T*^. Unfortunately, data that is represented in this way often heavily mixes the effect of different task parameters between latent dimensions. We would like to extract dimensions that dissociate our specific task conditions. To achieve this, demixed principal components analysis (dPCA) was performed^50^ using freely available code: http://github.com/machenslab/dPCA. In contrast to PCA, dPCA seeks to explain marginalized variance with respect to our specific task variables (stimulus and time), instead of merely explaining total variance. Unlike PCA, dPCA utilizes a separate encoder and decoder, such that the loss being optimized was ℒ = Σ _*ϕ*_ ℒ_*ϕ*_ = Σ _*ϕ*_ (∥X_*ϕ*_ − F_*ϕ*_ D_*ϕ*_ X^2^ ∥ + λ _*ϕ*_ ∥ F_*ϕ*_ D_*ϕ*_ ∥^2^), where *X*_*ϕ*_ is the marginalization of the full data with respect to each of our task parameters of interest and the *λ* term is a regularization parameter, preventing overfitting. Marginalizations of *X* can be obtained by averaging over all parameters which are not being investigated and subtracting all simpler marginalizations. In our case the marginalizations of interest were stimulus *x* time and time. The specific value of *λ* was determined using 5-fold cross-validation for each brain area, allowing each factor to have a different value of λ_*ϕ*_.

### Condition decoding analysis

To quantify auditory pattern information available on single trials, we trained multi-class linear decoders to classify the stimulus condition from population activity, separately for each recorded area and signal type (spikes vs. LFP). Activity was sampled every 100 ms between cue onset and 1 second after offset (8.3 seconds). For spikes, smoothed-spike trains were used and the smoothed activity from all units recorded in the area on a given session was concatenated across time to form a trial-level feature vector. For each area we concatenated data to form a design matrix (observations = trials; predictors = units x time for spikes and channels x frequency x time for LFP).

For LFPs, time samples across channels and frequency bands within the area were used analogously.

Within each area, data were de-meaned and reduced with PCA computed only on the training data. We conservatively retained the first M principal components, where M was equal to 20% of the number of trials, and projected both training and test data onto this subspace. Features were z-scored using statistics computed on the training set.

We used a 10-fold cross-validation scheme. On each fold, a multiclass error-correcting output-codes (ECOC) classifier with linear base learners was trained on the training trials and evaluated on the held-out fold. Base learners were logistic regression models with default L1 regularization. Decoder performance for each session/area was the fraction of correctly classified trials (sum of diagonal of the confusion matrix divided by its total). Confusion matrices were accumulated across sessions within each area for visualization.

To assess chance performance and set a conservative reference, we generated a permutation-based null by shuffling the condition labels and recomputing the fraction-correct 10,000 times. The reported “null level” for plotting was the 99.9th percentile of this distribution. Area-level performance was the mean across sessions, reported separately for spikes and LFP.

### Elapsed time prediction analysis

To test whether population activity carried a reproducible signal about trial progression independent of stimulus identity, we performed continuous time-regression from population activity to elapsed time within the trial. As above, activity was sampled every 100 ms from cue onset until 1 second after trial end (8.3 seconds). For spikes, smoothed firing rates were used. For each trial, the target vector was the elapsed time in seconds. For each area we concatenated data to form a design matrix (observations = time x trials; predictors = units for spikes and channels x frequency for LFP). As in the decoding analysis, we performed PCA on the training set, projected train/test data into that subspace, and standardized predictors using training-set statistics.

We fit an L1-regularized linear regression with 10-fold cross-validation. Performance was quantified as the Pearson correlation between predicted and true elapsed time computed across all held-out observations. For a chance benchmark, we constructed a null distribution by randomly permuting the target time series (10,000 permutations) and recomputing the correlation. The 99.9th percentile of this null served as the chance level. For each area and signal type (spikes, LFP), we report the mean cross-validated correlation across sessions, and also visualize the joint distribution of predictions and targets using 2-D histograms for interpretability.

### Stimulus encoding analysis

To quantify how strongly each neural population encoded the auditory tone sequence, we computed a cross-validated, noise-normalized Mahalanobis distance (“crossnobis distance”) between stimulus-weighted population activity patterns^51^. This approach measures the reliability of population representations associated with the tone waveform while accounting for shared noise covariance across neurons and time. Tone waveforms were smoothed using the same kernel as neural data as described above.

For this analysis, we used smoothed spike rates from 300ms after cue onset until trial end in 10 ms steps. This time window was smaller than previous analyses to prevent direct responses to reward in Experiment 1 and 3. Trials were divided into two independent splits (odd vs. even trials). Within each split, we constructed a stimulus regressor representing the instantaneous amplitude of the tone waveform over time, demeaned across trials within that split. Smoothed neural activity was multiplied by this regressor at each timepoint to generate a stimulus-weighted population vector, and these vectors were summed across all tone-active timepoints to yield one stimulus-weighted response vector per split. For the LFP version of this analysis, we concatenated frequency bands and channels to use as features (equivalent to neurons in the spiking analysis).

To normalize for correlated variability across neurons, we estimated the noise covariance matrix from trial-wise residuals (after subtracting trial means) and used this matrix to whiten both split-averaged vectors. The crossnobis distance was then computed as the dot product between the two whitened vectors. Because the distance is computed across splits, positive values indicate reliable stimulus-related structure rather than noise overfitting.

To assess statistical significance, we constructed a permutation-based null distribution by randomly shuffling trial labels of the stimulus regressor within each split 1000 times, recomputing the crossnobis distance each time. The observed value was z-scored relative to this null, yielding an encoding strength in units of standard deviations above chance. These z-scores were averaged across sessions to obtain a population-level measure for each brain area. This analysis provides a bias-free, cross-validated measure of how strongly neural population activity covaried with the timing of the auditory tones, independent of mean firing rate differences or overall activity level.

## Acknowledgements

Experiments were funded by: CIHR Operating Grants (Foundation Grant to J.A.P: 353197; Project Grant to J.A.P.: PJT-175010), Azrieli Foundation (via the Collaboration on Motor Planning, Execution and Resilience). S.N. was supported by a Connected Minds Postdoctoral Fellowship (CFREF). J.A.M. was supported by a VISTA research-enhanced position (CFREF), Banting Postdoctoral Fellowship, a Vector Institute Postgraduate Affiliation, and a BrainsCAN Postdoctoral Fellowship (CFREF). J.A.P. was supported by the Canada Research Chair program.

## Author Contributions

S.N., J.A.P., J.A.G., and J.A.M. conceived the study. S.N., M.K., R.K., J.A.P., and J.A.M. contributed to the monkey experiments. S.N. and J.A.M. contributed to the neural data analysis. All authors contributed to interpretation and writing. J.A.P. and J.A.M. contributed to the supervision and funding.

## Data availability

Data will be released in a public repository upon final publication.

## Code availability

Custom codes for data analysis were written in MATLAB and Python and are available from the corresponding author upon request.

## Supplementary Information

**Supplementary Table.** Information about each individual recording session.

| Session | Recorded neurons | Successful trials | Experiment |
| --- | --- | --- | --- |
| *May 27-31, 2024, 5 training sessions without neural recording* |  |  | 1 |
| 03-Jun-24 | SMA: 167, pre-SMA: 80 | 457 | 1 |
| 04-Jun-24 | PMd: 354, M1: 336 | 488 | 1 |
| 06-Jun-24 | MGB: 124 | 497 | 1 |
| 07-Jun-24 | S1: 131, GPi: 333 | 600 | 1 |
| *June 9-13, 2025, 5 training sessions without neural recording* |  |  | 2 |
| 09-Jun-25 | MGB: 177 | 419 | 2 |
| 10-Jun-25 | S1: 30 | 644 | 2 |
| 11-Jun-25 | M1: 196, PMd: 406 | 632 | 2 |
| 12-Jun-25 | pre-SMA: 107 | 659 | 2 |
| 13-Jun-25 | GPi: 591, SMA: 141 | 561 | 2 |
| *No training between Experiment 2 and 3* |  |  |  |
| 16-Jun-25 | GPi: 230 | 554 | 3 |

**Figure S1.**
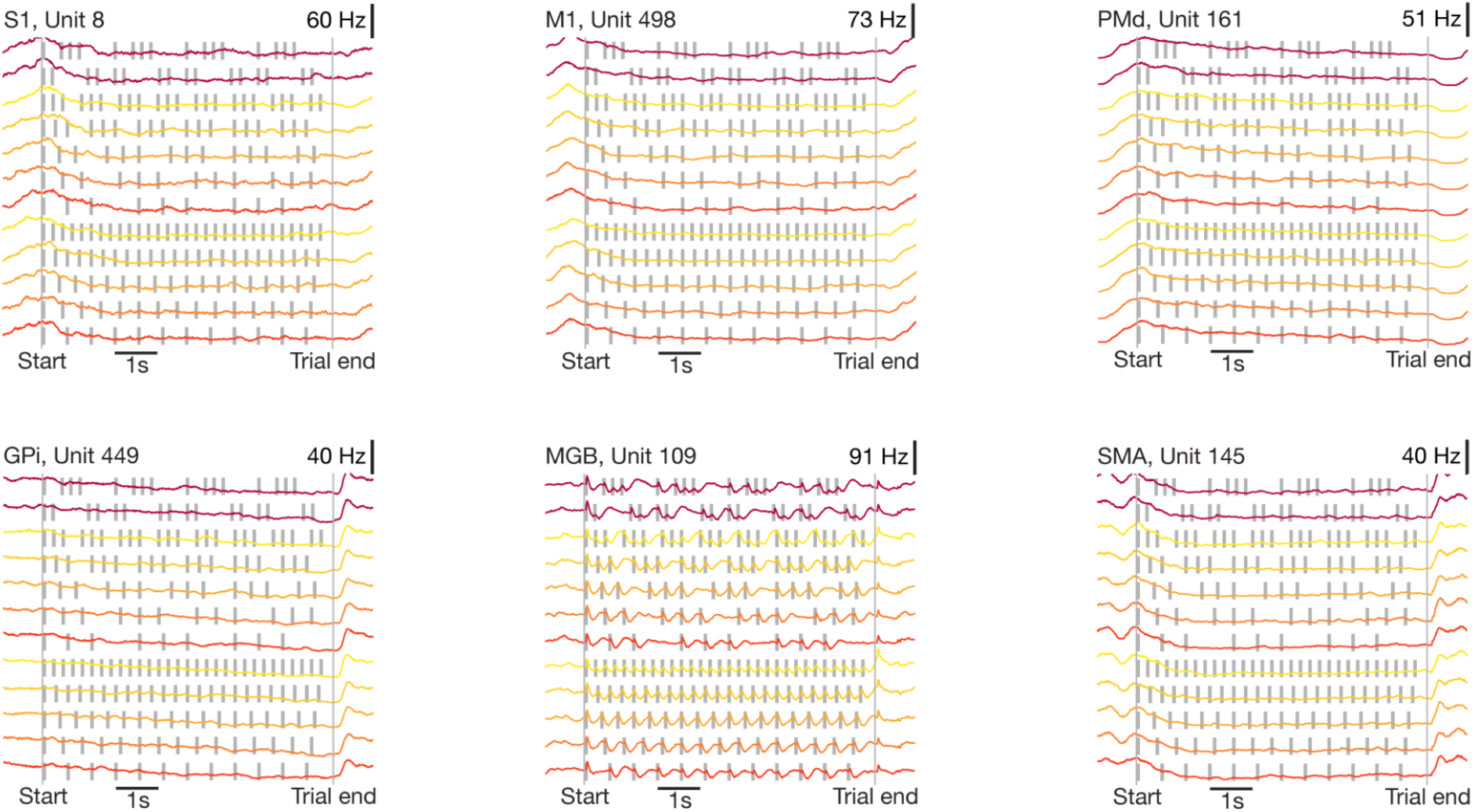
Single neuron examples from Experiment 1. The firing rates of example single neurons from six different brain regions show modulation to the temporal progression of trial, but only the MGB neuron shows a response to auditory tones (bottom, middle).

**Figure S2.**
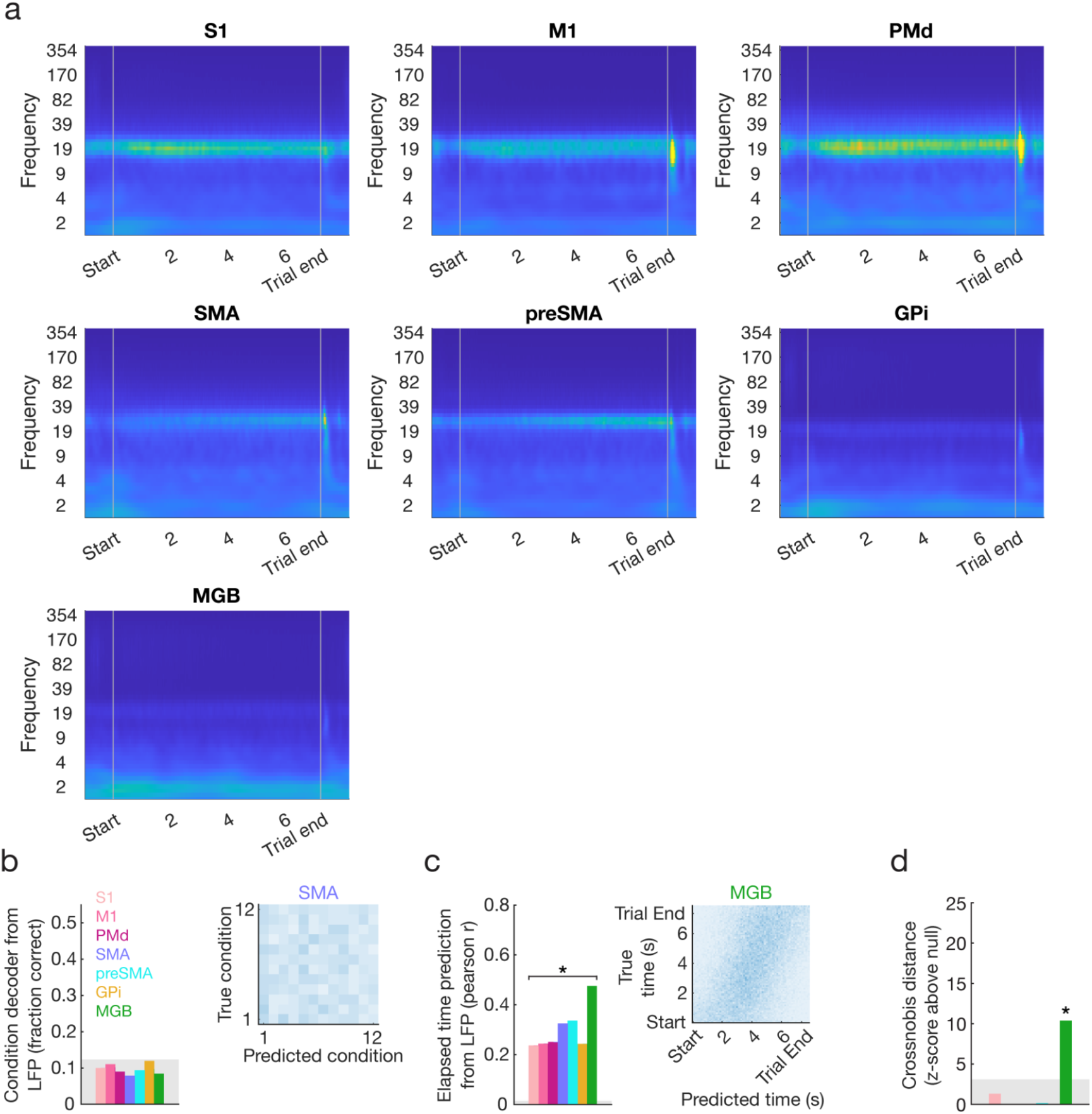
LFP of each area and decoding for Experiment 1. **a**, The LFP spectrogram was averaged across recording sites spanning seven different brain regions. **b, Left**, Average stimulus condition decoding accuracy across different brain areas. The shaded region denotes the chance level. **Right**, Accuracy of the condition decoder across all conditions in SMA. **c, Left**, Average elapsed time decoding accuracy across different brain areas. Star indicates p < 0.001 relative to chance level. **Right**, Accuracy of elapsed time decoder across all timepoints in MGB. **d**, Auditory pattern encoding quantified by crossnobis distance between odd-even splits of neural population activity weighted by normalized stimulus pattern. Stars indicate p < 0.001 relative to chance level.

**Figure S3.**
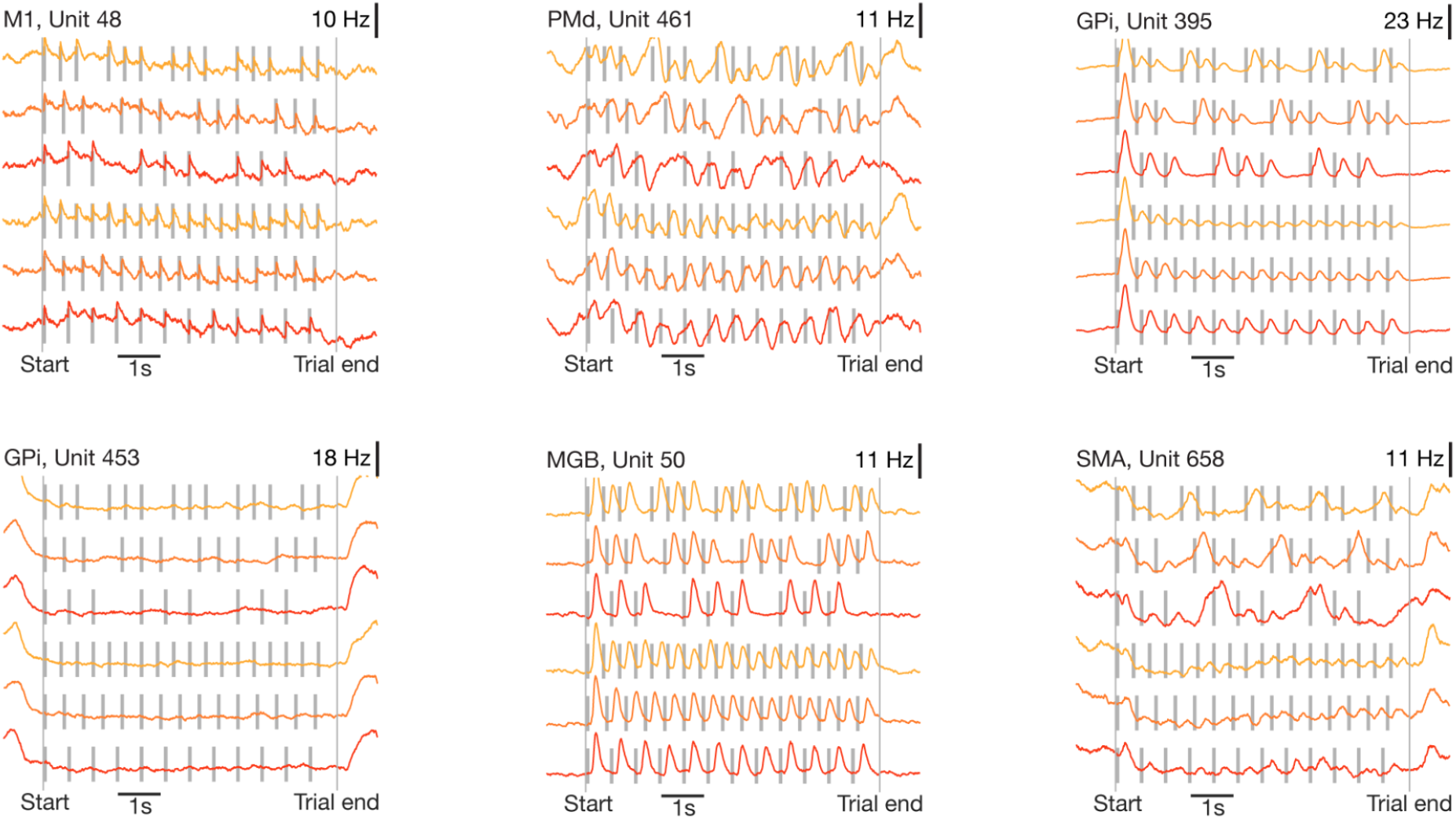
Single neuron examples from Experiment 2. The firing rates of example single neurons from six different brain regions show strong modulation to the auditory tones, which were paired with liquid reward in experiment 2 (bottom, middle).

**Figure S4.**
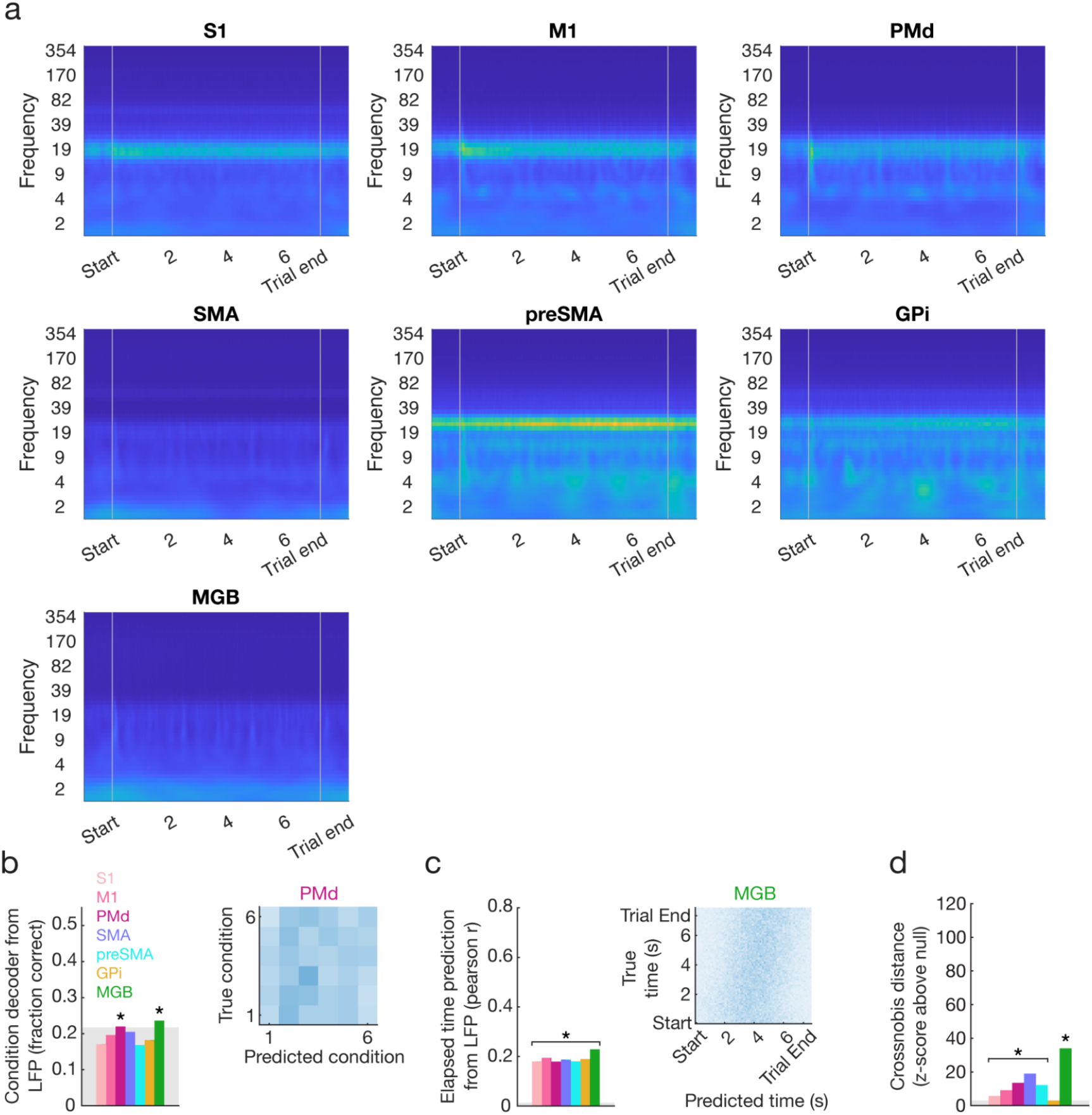
LFP of each area and decoding for Experiment 3. **a**, The LFP spectrogram was averaged across recording sites spanning seven different brain regions. **b, Left**, Average stimulus condition decoding accuracy across different brain areas. The shaded region denotes the chance level. **Right**, Accuracy of the condition decoder across all conditions in PMd. **c, Left**, Average elapsed time decoding accuracy across different brain areas. Star indicates p < 0.001 relative to chance level. **Right**, Accuracy of elapsed time decoder across all timepoints in MGB. **d**, Auditory pattern encoding quantified by crossnobis distance between odd-even splits of neural population activity weighted by normalized stimulus pattern. Stars indicate p < 0.001 relative to chance level.

**Figure S5.**
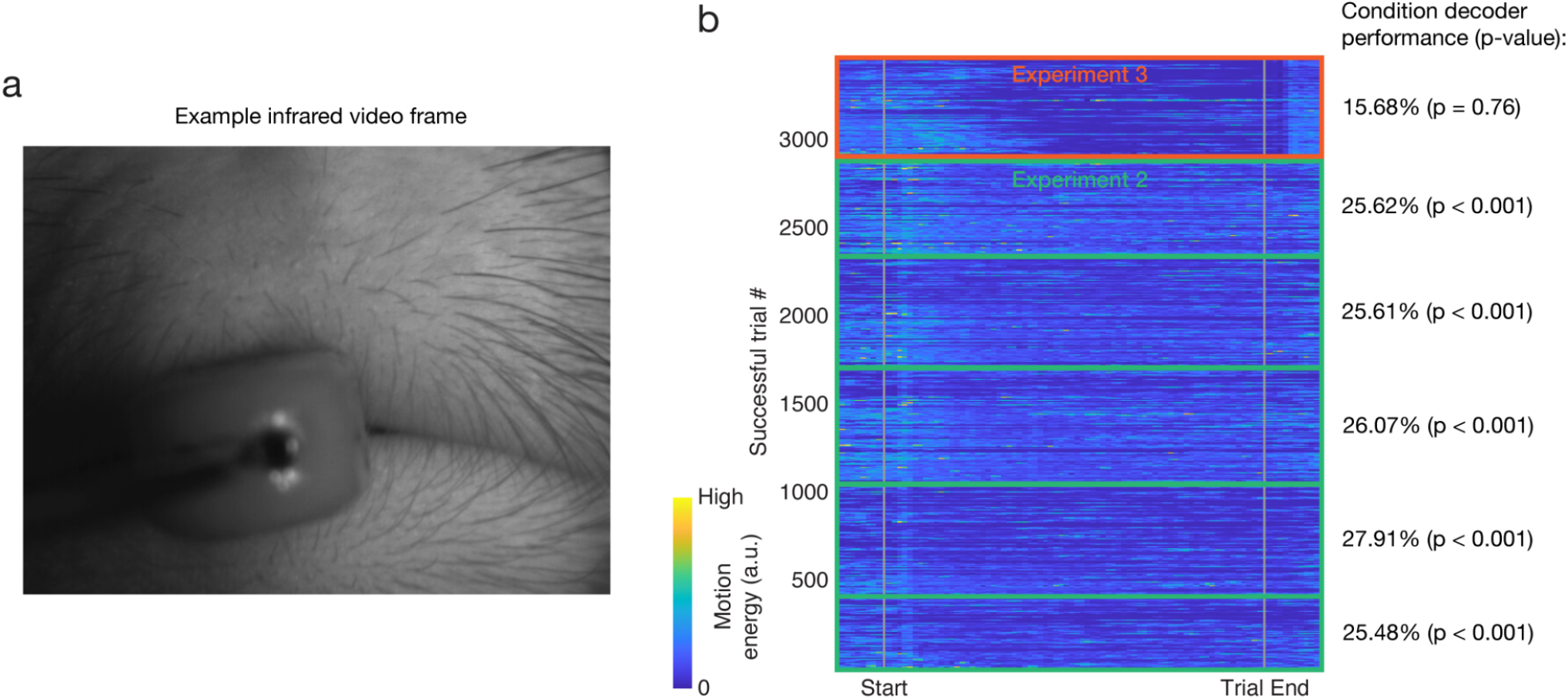
Face motion energy analysis. **a**, An example infrared video frame used for face motion energy analysis. **b**, Motion energy over time for all trials across Experiment 2 and 3. While it was possible to significantly decode which pattern of auditory stimulus was being played in Experiment 2, it was not possible in Experiment 3.

**Figure S6.**
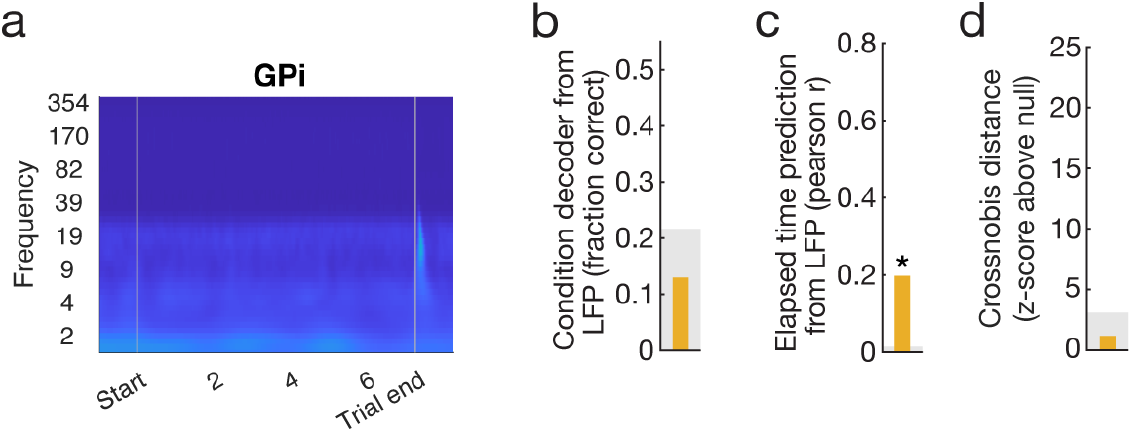
LFP of GPi area and decoding for Experiment 3. **a**, The LFP spectrogram was averaged across recording sites in GPi. **b**, Average stimulus condition decoding accuracy. The shaded region denotes the chance level. **c**, Average elapsed time decoding accuracy. Star indicates p < 0.001 relative to chance level. **d**, Auditory pattern encoding quantified by crossnobis distance between odd-even splits of neural population activity weighted by normalized stimulus pattern. Stars indicate p < 0.001 relative to chance level.

